# Motor Clustering Enhances Kinesin-driven Vesicle Transport

**DOI:** 10.1101/2024.10.23.619892

**Authors:** Rui Jiang, Qingzhou Feng, Daguan Nong, You Jung Kang, David Sept, William O. Hancock

## Abstract

Intracellular vesicles are typically transported by a small number of kinesin and dynein motors. However, the slow microtubule binding rate of kinesin-1 observed in *in vitro* biophysical studies suggests that long-range transport may require a high number of motors. To address the discrepancy in motor requirements between *in vivo* and *in vitro* studies, we reconstituted motility of 120-nm-diameter liposomes driven by multiple GFP-labeled kinesin-1 motors. Consistent with predictions based on previous binding rate measurements, we found that long-distance transport requires a high number of kinesin-1 motors. We hypothesized that this discrepancy from *in vivo* observations may arise from differences in motor organization and tested whether motor clustering can enhance transport efficiency using a DNA scaffold. Clustering just three motors improved liposome travel distances across a wide range of motor numbers. Our findings demonstrate that, independent of motor number, the arrangement of motors on a vesicle regulates transport distance, suggesting that differences in motor organization may explain the disparity between *in vivo* and *in vitro* motor requirements for long-range transport.

**Significance Statement:** Intracellular vesicles frequently travel long distances, despite having few kinesin and dynein motors. By reconstituting liposome motility with kinesin-1 motors, we demonstrate the need for high motor copy numbers for long-range transport when motors are randomly distributed on the liposome surface. We further show that motor clustering reduces the required motor number, emphasizing its potential role in enhancing transport efficiency. Our findings highlight the significance of motor organization in regulating intracellular transport and suggest that motor clustering, such as by scaffolding proteins or lipid domains, influences bidirectional transport outcomes.

## Introduction

Long-range intracellular transport relies on kinesin and dynein motors that travel towards microtubule plus- and minus-ends, respectively. Cargos typically possess both motor types, including diverse kinesin motors (1-3). While these cargos often travel tens of microns, surpassing individual motor run lengths, small vesicles (∼100 nm) generally have low motor counts: estimated to be one to three kinesin-1, two to six kinesin-2, and three to ten dynein motors (1, 2). A number of *in vitro* reconstitution studies using DNA scaffolds, optical tweezers, and giant vesicles have demonstrated that a small number of simultaneously engaged kinesin-1 motors (about three to ten) can achieve long-range transport (4-6); however, it remains unclear how simultaneous engagement of multiple motors is achieved and regulated. While it is generally assumed that most motors attached to a small cargo can easily access the microtubule (7, 8), the run length of 100-nm beads was found to only increase marginally when an estimated 73 kinesin-1 motors were bound (9). This high motor number requirement *in vitro* is further supported by the slow microtubule attachment rate of kinesin-1 (10, 11); however, it contrasts with the low motor copy number observed on purified intracellular vesicles. This discrepancy has generally been overlooked to date, as previous reconstitution studies using beads or vesicles did not experimentally quantify total motor copy numbers. Addressing this discrepancy requires a detailed characterization of the travel distance of vesicles containing a range of total motor copy numbers.

Although the robust motility by small numbers of motors in cells can be partly explained by microtubule-associated proteins (MAPs) like MAP7 recruiting motors to microtubules (12-14) and enhanced motor affinities due to microtubule post-translational modifications such as acetylation (15), the observation that purified endosomes travel long distances on MAP-free microtubules assembled *in vitro* (16) suggests that motor-cargo complexes employ additional regulatory mechanisms to achieve long-range transport. Increasing evidence suggests that motors may be clustered on the cargo membrane in cells. For instance, it has been shown that dynein clusters into cholesterol-rich lipid microdomains as phagosomes mature (8), and a scaffold protein for dynein-dynactin complexes, Septin 9, localizes to subdomains of the lysosomal membrane (17). Additionally, there is an emerging body of evidence that cargo adaptors can recruit two dynein motors or both kinesin and dynein motors, serving as another mechanism for motor clustering (18-21). While it has been demonstrated that dynein clustering into lipid microdomains is critical for the uninterrupted unidirectional retrograde movement of mature phagosomes (∼1 μm) (8), and clustering of monomeric Unc104 motors increases vesicle landing rate by promoting motor dimerization (22), it remains unknown whether motor clustering plays a role in regulating transport distance of small vesicles (∼100 nm) by processive dimeric kinesin motors.

To address these gaps, we reconstituted kinesin-1-driven vesicle (∼120 nm) transport *in vitro* to characterize transport distance as a function of total motor copy number and investigated whether motor clustering affects vesicle run length. Our data suggest that reconstituting long-range transport of small vesicles with randomly distributed kinesin-1 motors requires high motor copy number, while clustering motors into groups of three reduces the required motor number. These findings help to illuminate the discrepancy between motor counts *in vivo* and *in vitro*, underscoring the importance of motor organization in long-distance vesicle transport.

## Results and Discussion

### *In Vitro* Reconstitution of Liposome Transport by Kinesin-1 Motors

To reconstitute kinesin-driven vesicle motility *in vitro*, the truncated *Drosophila* kinesin-1 construct, K406GFP-SNAP, was attached to liposomes through a biotinylated DNA oligonucleotide (oligo) (Fig. 1A, inset, Supplementary Fig. 1). The mean diameter of liposomes was 123 ± 2 nm (95% CI of fit), measured by dynamic light scattering (DLS) (Fig. 1B). Upon conjugating biotinylated kinesin-1 motors to the liposomes, we removed unattached motors using a liposome flotation assay, as follows. The motor-liposome mixture was loaded to the bottom layer of a sucrose gradient, and following centrifugation, liposomes with attached motors floated to the top and were used for the motility assay, while unattached motors remained at the bottom (Supplementary Fig. 2). Removing unbound motors ensured that motor numbers on liposomes remained constant during the motility assay. Liposome motility on surface-immobilized microtubules was observed using total internal reflection fluorescence microscopy (TIRFM), while unlabeled microtubules were visualized by interference reflection microscopy (IRM) (23) integrated to the microscope (Fig. 1A, 1C, 2A) (24). To quantify total kinesin-1 copy numbers per liposome, we measured the integrated GFP fluorescence on liposomes immobilized on the microtubules using the non-hydrolysable ATP analog, AMPPNP, which locks the motors to the microtubule, thus localizing all motors within the 100-nm evanescent field of TIRFM (Fig. 1C). Motor copy number was then calculated by comparing the integrated GFP fluorescence intensity on liposomes to the fluorescence intensity distribution of single K406GFP-SNAP motors (Fig. 1D) (see Supplementary Information for details). This procedure was repeated at varying motor:vesicle ratios to achieve a range of motor copy numbers.

**Figure 1.**
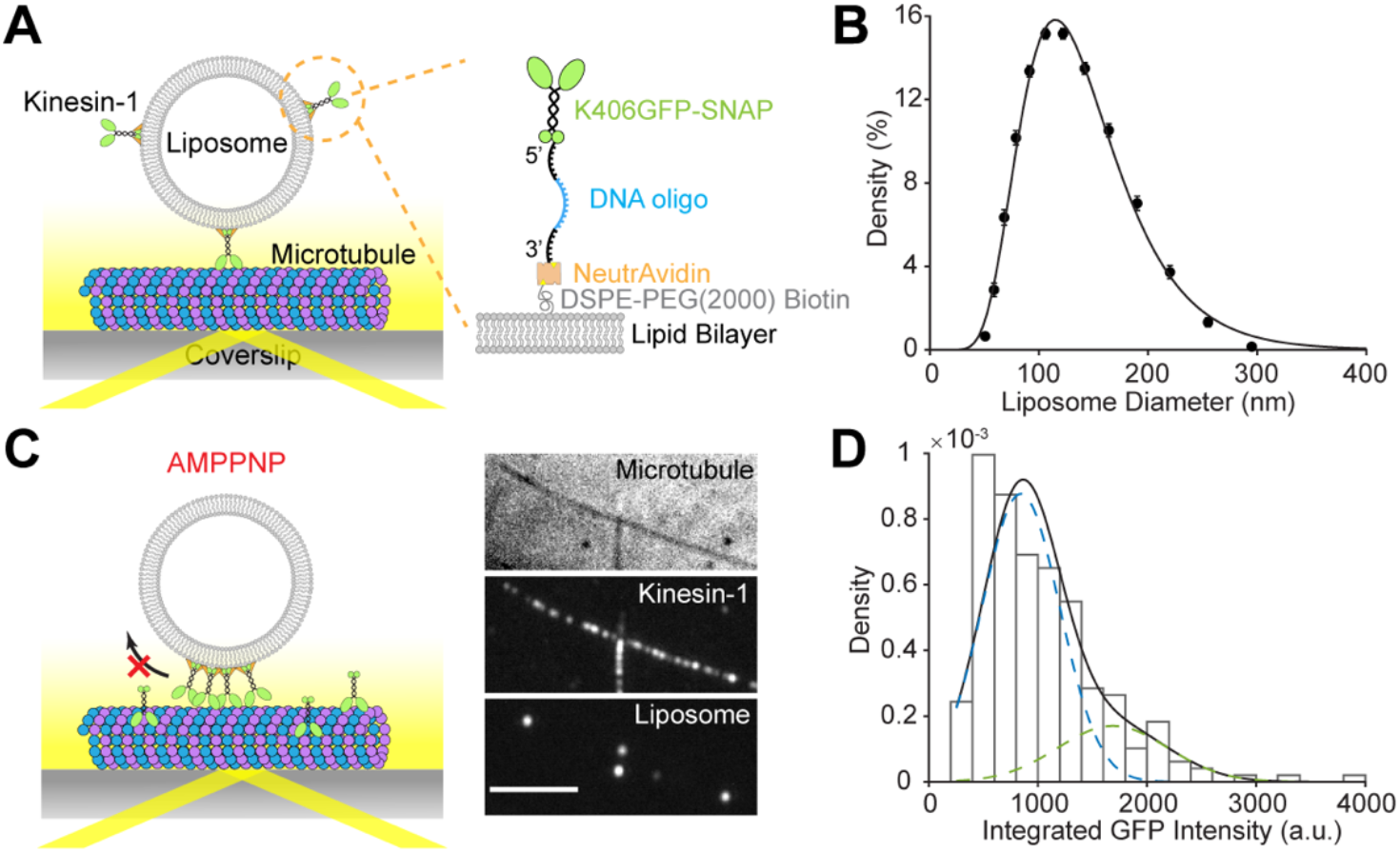
Reconstitution of liposome transport driven by kinesin-1. (A) Schematic of experimental setup. A truncated kinesin-1 motor (K406GFP-SNAP) is covalently linked to the amino group at the 5’-end of a DNA oligonucleotide (oligo) through BG-GLA-NHS. The biotin at the 3’-end of the DNA oligo attaches the motor to DSPE-PEG(2000)-Biotin on the liposome through NeutrAvidin. Liposome motility assay is carried out on surface-immobilized microtubules and imaged by total internal reflection fluorescence microscopy (TIRFM). (B) Vesicle size distribution measured by dynamic light scattering (DLS). Black circles show data averaged from three samples. Error bars indicate SEM. Black line indicates fit to a lognormal distribution, weighted by inverse of SEM. Fit parameters (95% CI of fit): μ = 123 ± 2 nm, and σ = 48 ± 2 nm. (C) Snapshot showing co-localization of kinesin-1 motors and liposomes on the microtubules in AMPPMP. Scale bar, 5 μm. (D) Example histogram of integrated GFP intensity of single kinesin-1 motors (N = 246). Black line shows fit to a sum of two normal distributions, where the mean and variance of the second peak (2μ, 2σ^2^) is set to be twice of those of the first peak (μ, σ^2^). Blue dashed line shows the contribution of the first peak and green dashed line shows contribution of the second peak. Fit parameters (±95% CI of fit): μ = 843 ± 77, σ = 357 ± 49, amplitude of the first peak A_1_ = 0.78 ± 0.11.

### Long-Range Liposome Transport Requires High Kinesin-1 Copy Numbers

Single K406GFP-SNAP motors exhibited a velocity of 623 ± 123 nm/s (mean ± SD, N = 406) and a run length of 0.49 ± 0.02 μm (95% CI of fit) (Supplementary Fig. 3). Liposomes moved at a similar but slightly slower speed than single kinesin-1 motors (Fig. 2A, Supplementary Fig. 5), which was not unexpected since the average speed of liposomes would be more strongly impacted by the slower motors in the group. This slowing contrasts with previous findings that coupling of kinesin-1 and myosin Va to cargo through a fluid lipid membrane increased transport velocities (25, 26). The larger cargo size in the previous kinesin-1 studies (500 nm vs. 100 nm here) may play a role because one potential mechanism for faster velocities is that when slower trailing motors detach, the cargo centers around the remaining motors. This recoil causes a larger positive displacement for larger cargo. For myosin Va, different load-dependent detachment kinetics from kinesin-1 could also contribute to the faster velocities.

**Figure 2.**
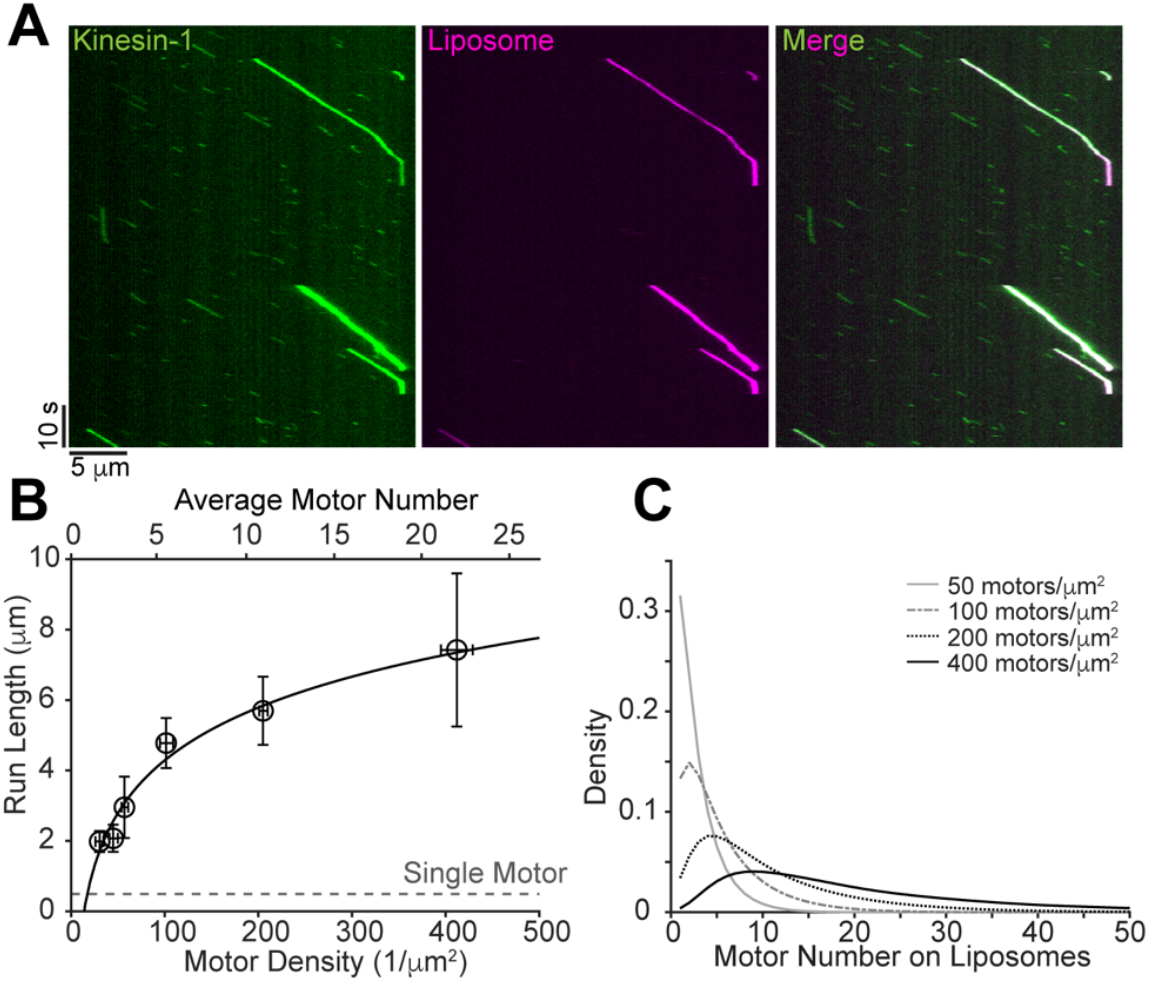
Liposome run lengths increase with higher kinesin-1 copy numbers. (A) Example kymographs showing liposome motility driven by multiple kinesin-1 motors in ATP. (B) Liposome run lengths at varying surface motor densities/numbers. Open black circles represent the mean of raw data, vertical error bars represent 95% CI and horizontal error bars are calculated by a 10% increase in the sum of squared residuals. Black line indicates fit to a logarithmic function: y = a·log_10_x + b. Fit parameters: a = 5.0 ± 1.5, b = −5.7 ± 2.7. Gray dashed line indicates single motor run length. (C) Motor number distribution on liposomes at varying motor densities, according to Supplementary Information Eq. 10.

The run length of liposomes driven by multiple kinesin-1 motors increased in a logarithmic manner with motor surface density - the fitting indicated that a 10 μm increase in run length requires a 100-fold increase in motor density (Fig. 2B). To facilitate data interpretation, we converted the motor surface density to motor number distribution using Supplementary Information Eq. 10 (Fig. 2C); the corresponding mean values of these distributions are included as a second x-axis at the top of Fig. 2B. Our measurement suggests that to achieve a run length of 10 μm requires a kinesin-1 surface density of 1000/μm^2^, which equates to ∼30 motors for a 100 nm vesicle. This measurement is in good agreement with the previous prediction that ∼35 kinesin-1 motors are needed to transport a 100-nm vesicle for 10 μm based on slow kinesin-1 attachment rate measured on supported lipid bilayers (10). This result demonstrates that long-range vesicle transport requires high motor copy numbers when the motors are randomly distributed on the vesicle surface. Our results contrast with both the low motor copy number on purified intracellular vesicles (1, 2), and the presumption that most motors attached to such small cargos can simultaneously interact with the microtubule (7, 8).

### Induction of Motor Clustering by A DNA Scaffold

One possible reason that our reconstitution assay does not recapitulate the long-range transport with small motor numbers observed *in vivo* is that motors may not be randomly distributed on the cargo in cells and the ability of motors to coordinate their behavior depends strongly on the distances between the motors. To test this possibility, we investigated whether motor clustering on the liposome surface can enhance transport efficiency by using a DNA scaffold to precisely cluster motors into groups of three. The clustering oligo contains three repeats of a sequence complementary to a portion of the motor oligo, enabling it to induce clustering of up to three kinesin-1 motors (Fig. 3A). Electrophoretic mobility shift assays (EMSA) confirmed cluster formation, showing a molecular weight shift corresponding to a cluster of three motors at high motor:clustering oligo ratios and lower molecular weight bands appearing at increasing clustering oligo concentrations (Supplementary Fig. 4). To ensure that the introduction of the clustering oligo only changes the organization of existing motors on the liposomes rather than recruiting additional motors, we introduced the clustering oligo after excess motors were removed by the liposome flotation assay. To distinguish liposomes with clustered motors from those without, we labeled the clustering oligo with Cy3 in the motility assay. Cy3+ liposomes indicated the presence of motor clusters (Fig. 3B, 3C), while liposomes without clusters were Cy3-. The Cy3 labeling also allowed us to measure the number of motors per liposome with and without clusters separately, to ensure the effects of motor clustering were compared between liposomes with similar motor counts.

**Figure 3.**
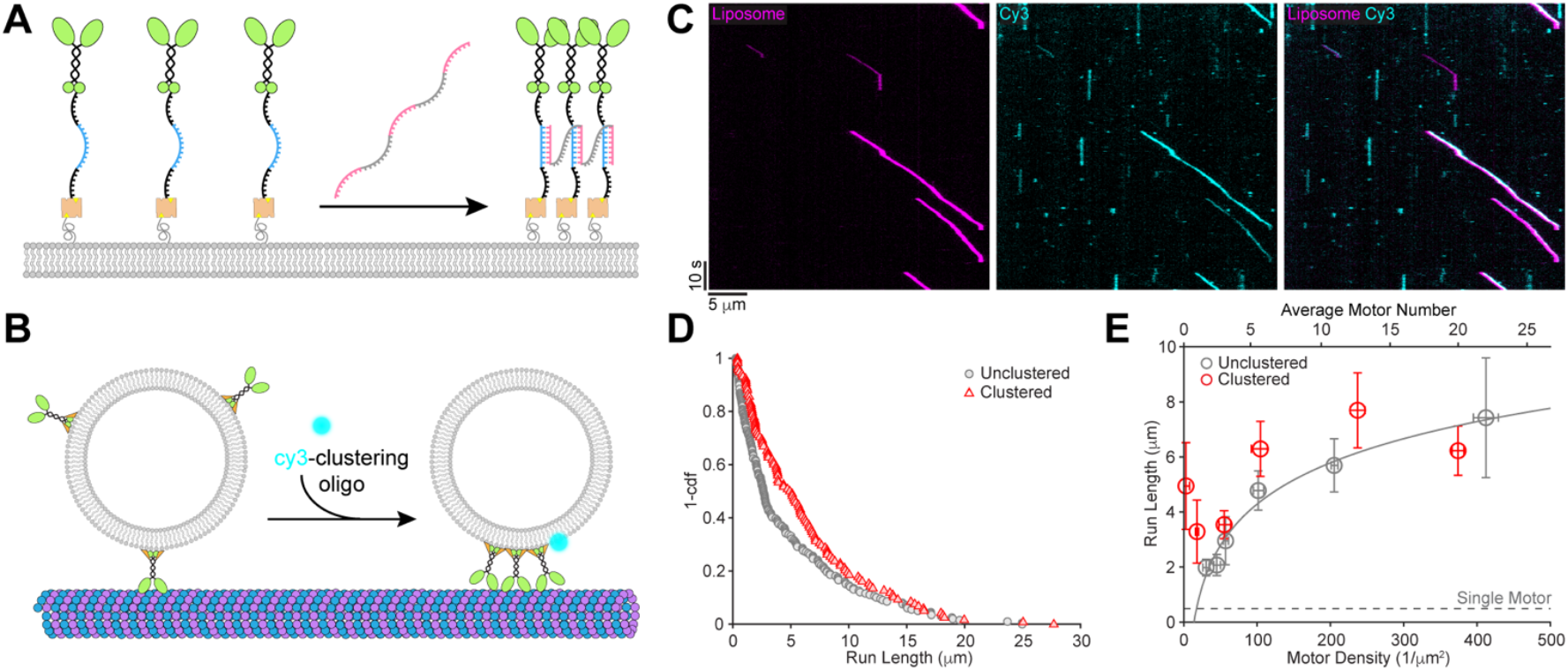
Motor clustering increases liposome travel distances. (A) Schematic of the clustering scaffold design. The clustering oligo contains three repeats of a sequence (pink) complementary to a portion (blue) of the motor oligo. Addition of the clustering oligo will induce formation of clusters of up to three motors (Supplementary Fig. 4). (B) Schematic of vesicle motility assay with clustered motors. The clustering oligo is labeled with Cy3. Liposomes co-localized with Cy3 signal contain cluster(s) of up to three motors. (C) Example kymograph showing motility of liposomes co-localized with Cy3-clustering oligo. (D) Liposome run length distributions at ∼100 motors/μm^2^ when motors are unclustered (gray) or clustered (red). (E) Summary of liposome run lengths at varying motor densities when motors are unclustered (gray, same as Fig. 2B) or clustered (red). Gray dashed line indicates single motor run length. Unclustered liposomes included both Cy3-liposomes and those from assays in which the clustering oligo was omitted.

### Motor Clustering Improves Liposome Travel Distances

The most notable result from this study is that clustering of just three kinesin-1 motors strongly boosted liposome transport distance across various motor densities without affecting transport speed (Fig. 3D, 3E, Supplementary Fig. 5). At very low motor densities (< 20 /μm^2^), where only a small fraction of liposomes had motors associated, we rarely observed motility without clustering, while liposomes with clustered motors showed a 6-to 10-fold increase in run length compared to single motors, likely representing transport by a single motor cluster. At moderate motor densities (100 – 200 /μm^2^), where each liposome had 6-10 motors on average, clustering still substantially extended run length. However, at high motor densities (> 400 /μm^2^), clustering did not affect run length. It is possible that the effects of clustering were masked by the microtubule length limits, as we frequently observed runs that reached the end of a microtubule (Fig. 3C). Additionally, the advantage conferred by clustering could be diminished at high densities because clustering reduces the effective motor unit density (defined as a single unclustered motor or a single cluster of motors) by three fold and therefore reduces the probability that a second motor cluster will engage with the microtubule. While the motor clusters will diffuse more slowly than single motors, we do not expect that this slower diffusion will alter motor attachment rate in our assay. Analogous to our clustering scenario, previous work showed that having three lipid-binding PH domains reduced the single-molecule diffusion coefficient by three-fold compared to proteins with a single PH domain (27). Importantly, in previous work the attachment rate of membrane-bound kinesin-1 was shown to be unchanged by a four-fold reduction in membrane diffusivity (10). In conclusion, our finding that motor clustering enhances vesicle transport distance driven by small motor number suggests that motor clustering can serve as an important regulatory mechanism in cells, particularly for motors with a slow attachment rate and those that are not superprocessive. This clustering mechanism could in principle be a regulator of the net directionality of bidirectional transport of vesicles by teams of kinesin and dynein motors. The present study further motivates investigations into the molecular mechanisms driving motor cluster formation on membranous cargos in cells.

## Materials and Methods

### Protein Construct and Purification

The K406GFP-SNAP construct contained the first 406 aa of *Drosophila* kinesin heavy chain fused to an eGFP followed by a SNAP-tag and a 6×His-tag at the C-terminal. Molecular biology was carried out using Gibson Assembly and Q5 site-directed mutagenesis kit (New England Biolabs, NEB). The protein was expressed in Tuner (DE3) cells (Sigma), and purified by affinity chromatography (28). Motor concentration was measured by the absorbance or fluorescence of GFP at 488 nm.

### Conjugation of Kinesin-1 to DNA Oligo

The motor oligo (Integrated DNA Technologies (IDT)) was modified by an amino group at the 5’-end and a biotin at the 3’-end (5’-/5AmMC6/GTCAATAATACGATAGAGATGGCAGAAGGGAGAGGAGTAGTGGAGGTAGAGTCAGGGCGAGAT/3Bio/-3’). The oligo was reacted with 20× excess of BG-GLA-NHS (NEB) in 100 mM sodium borate (Alfa Aesar) containing 50% (v/v) DMSO at 21 °C for 30 min. The excess BG-GLA-NHS was removed by desalting the reacted mixture into 1× PBS + 1 mM DTT + 1 mM MgCl_2_ using a PD MiniTrap desalting column with Sephadex G-25 resin (Cytiva). Oligo concentration was measured using a Nanodrop, and successful conjugation of BG-GLA-NHS was confirmed by running the sample on a 10% TBE-Urea gel (Invitrogen). The labeled oligo was mixed with K406GFP-SNAP (dimer) at a ratio of 1.5:1 and incubated on ice for 1 h. Excess oligo was removed by running the sample through a Ni column. Single oligo-labeling of motors was confirmed by running the sample on a 6% Tris-glycine gel (Invitrogen) under native conditions. Motors were stored in elution buffer supplemented with 10 μM MgATP, 1 mM DTT, and 10% sucrose at −80 °C.

### Electrophoretic Mobility Shift Assay (EMSA)

Oligo-labeled K406GFP-SNAP was mixed with the clustering oligo (5’-CCACTACTCCTCTCCCTTCTGCCCAACCACGACAAATCCCACTACTCCTCTCCCTTCTGCCCAACCACGACAAATCCCACTACTCCTCTCCCTTCTGCC-3’, IDT) in EMSA sample buffer (62.5 mM Tris·HCl, pH 7.5, 2% SDS, 10% glycerol, 0.005% Bromophenol Blue, 2.5% 2-β Mervaptoethanol), heated at 90 °C for 5 min, and left at RT for 30 min to allow the oligos to anneal. The samples were then run on a 6% Tris-glycine gel under denaturing condition at 100 V on ice. The gel was imaged by Coomassie staining.

### Liposome Preparation

1-palmitoyl-2-oleoyl-glycero-3-phosphocholine (POPC), 1,2-disteroyl-*sn*-glycero-3-phosphoethanolamine-N-[biotinyl(polyethylene glycol)-2000] (DSPE-PEG(2000)-Biotin) were purchased from Avanti. 1,2-dioleoyl-*sn*-glycero-3-phosphoethanolamine labeled with Atto 647N (Atto 647N DOPE) was purchased from Sigma. Lipids were dissolved in chloroform and POPC:DSPE-PEG(2000)-Biotin:Atto 647N DOPE were mixed at 99:1:0.05. The chloroform was evaporated under vacuum, and the lipids were rehydrated with BRB80 (80 mM Pipes, 1 mM EGTA, 1 mM MgCl_2_; pH 6.9) at 10 mg/ml. The lipid suspension was subjected to 10 freeze-thaw cycles in liquid nitrogen and a warm water bath. Liposomes were formed by extruding the lipids 21 times through a stack of two 100-nm membrane filters using a mini-extruder (Avanti). Liposomes were stored at 4 °C and used within a week.

### Coverslip Silanization

Glass coverslips (#1.5, 18×18 mm, Corning) were cleaned in hot 7x detergent (MP Biomedicals) diluted 1:7 with ultrapure water for 2 h. Coverslips were then rinsed thoroughly with ultrapure water, blown dry with nitrogen, and cleaned by a plasma cleaner (Harrick Plasma). 1H,1H,2H,2H-Perfluorodecyltrichlorosilane (Alfa Aesar) was allowed to deposit onto the coverslips in a vacuum desiccator overnight.

### Purification of Active Kinesin-1 by Microtubule Pelleting Assay

Taxol-stabilized microtubules were polymerized from bovine brain tubulin, as described previously (28). Oligo labeled K406GFP-SNAP (400 nM) was incubated with 4 μM NeutrAvidin (Thermo Scientific) in BRB80 containing 10 μM taxol, 500 μM MgAMPPNP, 1 mM DTT, 1 mg/ml casein and 1 mg/ml BSA for 5 min at RT. Microtubules (1.5 μM) were added to the mixture and incubated for another 5 min. The microtubules with attached motors were pelleted by centrifuging in an Airfuge (Beckman Coulter) at 20 psi for 3 min. Inactive motors and excess NeutrAvidin in the supernatant were discarded. The pellet was resuspended in BRB80 supplemented with 10 μM taxol, 2 mM MgATP, 1 mM DTT, 1 mg/ml casein, 1 mg/ml BSA and incubated for 10 min at RT to release the active motors. A second spin in the Airfuge at 30 psi for 5 min removed microtubules with irreversibly bound motors. The active motor-NeutrAvidin complexes in the supernatant was saved and kept on ice.

### Liposome Flotation Assay

The active motor-NeutrAvidin complexes purified by microtubule pelleting assay were incubated with liposomes in flotation buffer (100 μM MgATP, 1 mM MgCl_2_, 1 mM DTT, 1 mg/ml casein and 1 mg/ml BSA in BRB80) and incubated on ice for 30 min. The sample was mixed with equal volume of flotation buffer containing 60% (w/v) sucrose to form the bottom layer of the gradient. The middle and top layers contained flotation buffer with 25% and 0% sucrose, respectively. The samples were centrifuged in a Type 50.2 Ti rotor at 45,000 rpm, 4 °C for 45 min in an ultracentrifuge (Beckman Coulter). The top fraction containing liposomes and the attached motors was harvested after the flotation assay and motor concentration was measured by GFP fluorescence.

### Motor Number Quantification and Motility Assay

A narrow flow cell was assembled by sandwiching two strips of double-sided tape (∼2 mm apart) between a salinized coverslip and a glass slide. Anti-tubulin antibody (Sigma, 1:50 diluted in BRB80) was incubated in the chamber for 5 min, followed by a wash with BRB80 and incubation with 5% Pluronic F-127 in BRB80 for 5 min. The chamber was washed with BRB80. To align the microtubules on the surface, ∼5× chamber volumes of unlabeled taxol-stabilized microtubules (80 nM) were quickly flowed through the chamber and incubated for 30 s. This step was repeated once, followed by a wash with BRB80 + 10 μM taxol. To visualize liposome motility, liposomes were diluted in motility buffer (10 μM taxol, 2 mM MgATP, 10 mM DTT, 1 mg/ml casein, 1 mg/ml BSA, 20 mM D-glucose, 0.02 mg/ml glucose oxidase, 0.008 mg/ml catalase in BRB80) and imaged at 5 fps. To quantify motor copy numbers, liposomes were diluted in landing buffer (10 μM taxol, 1 mM MgAMPPNP, 10 mM DTT, 1 mg/ml casein, 1 mg/ml BSA, 20 mM D-glucose, 0.02 mg/ml glucose oxidase, 0.008 mg/ml catalase in BRB80) and allowed to land on the microtubules by incubating for 5 min. Excess liposomes were removed by a wash with landing buffer before imaging. To induce motor clustering, 5 nM of clustering oligo labeled with Cy3 at the 3’-end (IDT) was incubated with the liposomes on ice for 30 min before conducting motility or landing assays. Imaging was carried out on an integrated multi-wavelength microscope combining TIRFM and IRM modalities (24).

### Data Analysis

Liposome and single K406GFP-SNAP motility was analyzed by kymographs using Fiji (29). Single K406GFP-SNAP run length was fit to a single exponential distribution, where runs shorter than 0.22 μm (3 pixels) were eliminated from the fitting. Mean liposome travel distance was calculated by bootstrapping. Motor number on liposomes was measured by ratiometric comparison (30) of GFP fluorescence from K406GFP-SNAP in AMPPNP, as follows. GFP intensity was measured on a mean projection across 10 frames to increase the signal-to-noise ratio. The fluorescence intensity of single motors or multiple motors on a liposome was determined by subtracting the raw integrated intensity by the background intensity averaged from two adjacent regions of the same size. The fluorescence intensity of single K406GFP-SNAP was fitted to a sum of two normal distributions, where the mean and variance of the 2^nd^ peak were set to be twice of those of the 1^st^ peak. The mean of the 1^st^ peak corresponded to the signal from one GFP and the probability of a GFP being fluorescent was calculated based on the amplitude of the first peak. The expected GFP intensity distribution on liposomes with known motor densities were then computed based on the intensity distribution of single GFPs, the probability of a GFP being fluorescent, and the distribution of liposome sizes (See Supplementary Information for details). The measured GFP intensity distribution at each motor/liposome mixing ratio was then compared to the expected distributions, and motor density was determined by minimizing the sum of squared residuals from the expected distribution. Error in motor density estimation was determined by a 10% increase in the summed squared residuals. All fitting was carried out using MATLAB (The MathWorks). The motor number distribution on liposomes were calculated according to Supplementary Information Eq. 10.

## Data Availability

The authors declare that the data used to generate all plots in this study are available in the source data file. The experimental tracks are available from the corresponding author upon request. Source data are provided in this paper.

## Code Availability

The MATLAB scripts for data processing are available from the corresponding author upon reasonable request.

## Acknowledgements

We are grateful to Adheshwari Ramesh for her help with molecular biology of the K406GFP-SNAP construct, Ekaterina Bazilevskaya at the Penn State Materials Characterization Lab for her help with the dynamic light scattering (DLS) measurement, members of the Hancock laboratory for helpful discussions. This work was funded by NIH Grant R35GM139568. R.J. was supported by NIH Grant T32 GM108563 and D.S. was supported by R01GM136822.

## Author Contributions

R.J., Q.F., and W.O.H. designed research; R.J. and Y.J.K. performed research; D.N. constructed the microscope; R.J., Y.J.K., D.S., and W.O.H. analyzed data; R.J. and W.O.H. wrote the paper.

## Competing Interests

The authors declare no competing interests.

## Supplementary Information

### Supplementary Text

#### Distribution of integrated GFP intensity of single kinesin-1 motors

Each kinesin-1 dimer has two GFPs and the probability of each GFP being fluorescent (termed p) is independent. Since only motors with a fluorescent signal were included in the measurement, the resultant intensity distribution can be described by a sum of two independent normal distributions.

The integrated fluorescent intensity of a single GFP follows a normal distribution N(μ,σ^2^), with a probability density function (pdf):

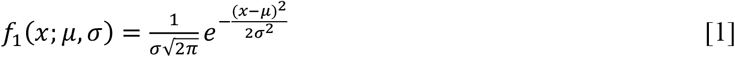

The integrated fluorescent intensity of two GFPs follows a normal distribution N(2μ,2σ^2^) with a pdf:

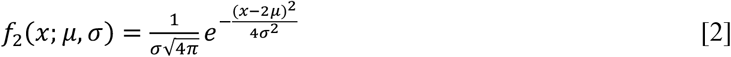

The amplitude of f_1_ (termed A_1_) can be calculated from p:

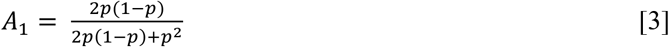

The pdf of integrated GFP intensity of a single kinesin-1 motor is therefore:

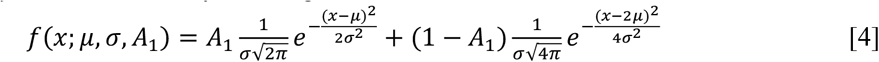

On each experimental day, the GFP intensity of single kinesin-1 motors were measured and fit to Eq. **4** by maximum likelihood estimation (MLE) to obtain μ and σ (Fig. 1D).

#### Distribution of integrated GFP intensity of n motors

The probability of *i* GFPs among the *n* motors being fluorescent:

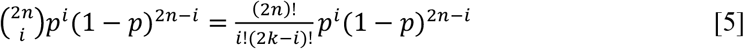

Since liposomes without any GFP signal were not included in the analysis, the probability calculated in Eq. **5** is normalized by the probability of any number of GFPs being fluorescent:

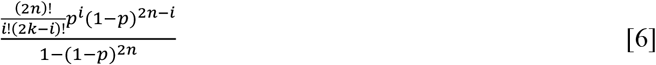

The integrated intensity of *i* GFPs follows a normal distribution N(iμ,iσ^2^), with a pdf:

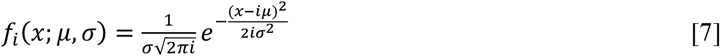

The pdf of the integrated GFP intensity of *n* motors is calculated as:

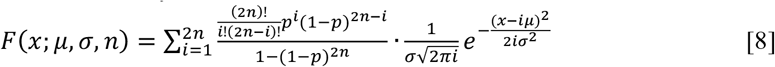

#### Distribution of motor numbers on liposomes

According to the liposome diameter distribution measured by DLS (Fig. 1B), the probability that a liposome is of area A_j_ (termed p_Aj_) is calculated (j = 1-13). Assuming that the surface motor density (ρ) is the same across all liposome sizes under each experimental condition and that motor number on liposomes of area A_j_ follows a poisson distribution Pois(ρA_j_), the pdf of motor number across all liposome sizes can be calculated as:

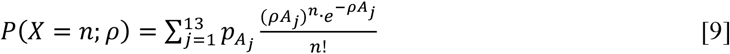

Since vesicles with zero motors were excluded from the analysis, the pdf is normalized as:

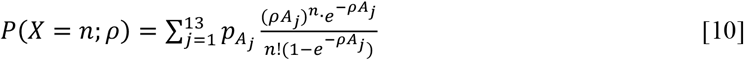

#### Distribution of integrated GFP intensity on liposomes

Combining Eqs. **8** and **10**, the pdf of the integrated GFP intensity on liposomes can be expressed as:

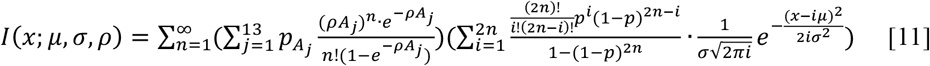

## Supplementary Figures

**Supplementary Figure 1.**
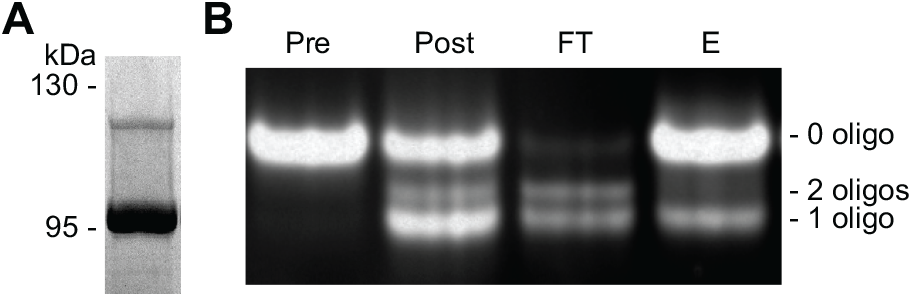
Characterization of K406GFP-SNAP by electrophoresis. (A)Coomassie staining of SDS-PAGE showing unlabeled K406GFP-SNAP monomer at 93 kDa, and K406GFP-SNAP-oligo monomer at 114 kDa. (B) Native PAGE showing K406GFP-SNAP samples at different stages of the oligo labeling process, imaged by GFP fluorescence. Pre, K406GFP-SNAP dimers before labeling. Post, K406GFP-SNAP and oligo mixture after the reaction. FT, the portion of the post sample that does not bind to Ni resin (flow through) in the second round of purification. E, eluted sample from Ni resin. Note that the running speed of the 3 species on a native gel is K406GFP-SNAP dimer + 1 oligo > K406GFP-SNAP dimer + 2 oligos > K406GFP-SNAP dimer. This is supported by the preferential exclusion of the two-oligo population from the Ni resin.

**Supplementary Figure 2.**
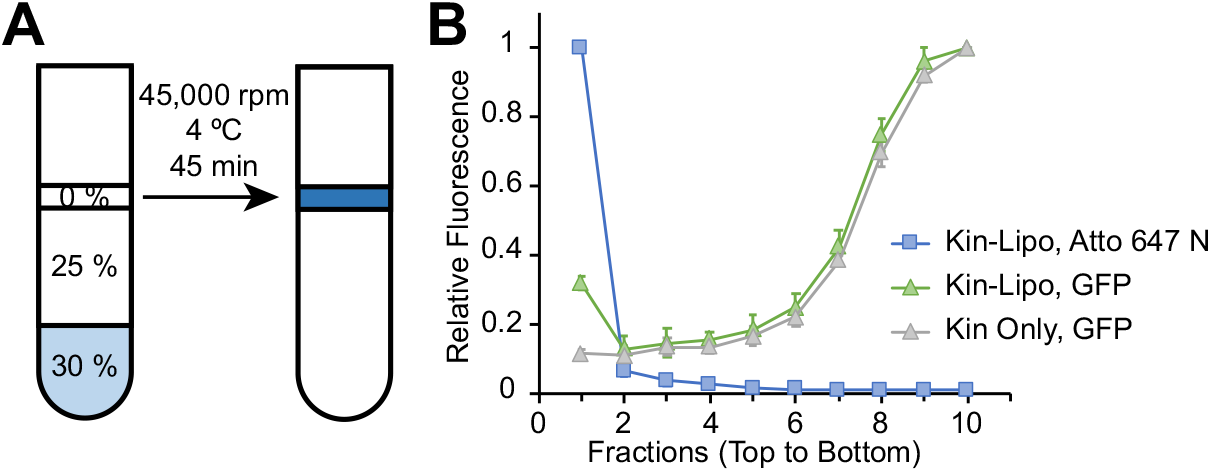
Liposome flotation assay. (A) Schematic of liposome flotation assay. Liposome and motor mixture is loaded to the bottom layer of a sucrose gradient. Upon centrifugation, liposomes with the attached motors ‘float’ to the top layer, while the unattached motors remain at the bottom. (B) Sample distribution post flotation. Blue squares and green triangles show relative fluorescence signal of liposomes and motors, respectively, when the mixture is loaded to the bottom layer prior to flotation. Grey triangles show motor distribution post flotation when liposomes are omitted. Each data point represents the mean of three independent measurements. Error bars indicate SD.

**Supplementary Figure 3.**
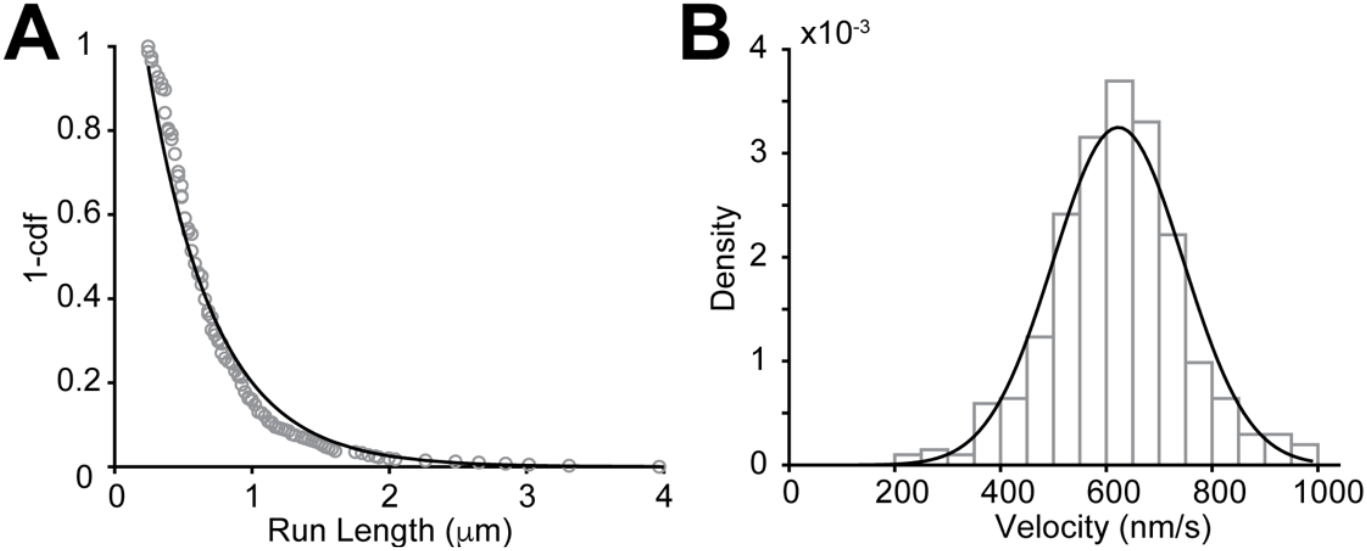
Single molecule characterization of K406GFP-SNAP. (A) Single molecule run length of K406GFP-SNAP. Grey open circles show distribution of raw data (N = 406), black line shows exponential fitting, λ = 0.49 ± 0.02 μm (95% CI of fit). (B) Histogram of K406GFP-SNAP single molecule velocity (N = 406) and fit to a normal distribution. v = 623 ± 123 nm/s (mean ± SD from fit).

**Supplementary Figure 4.**
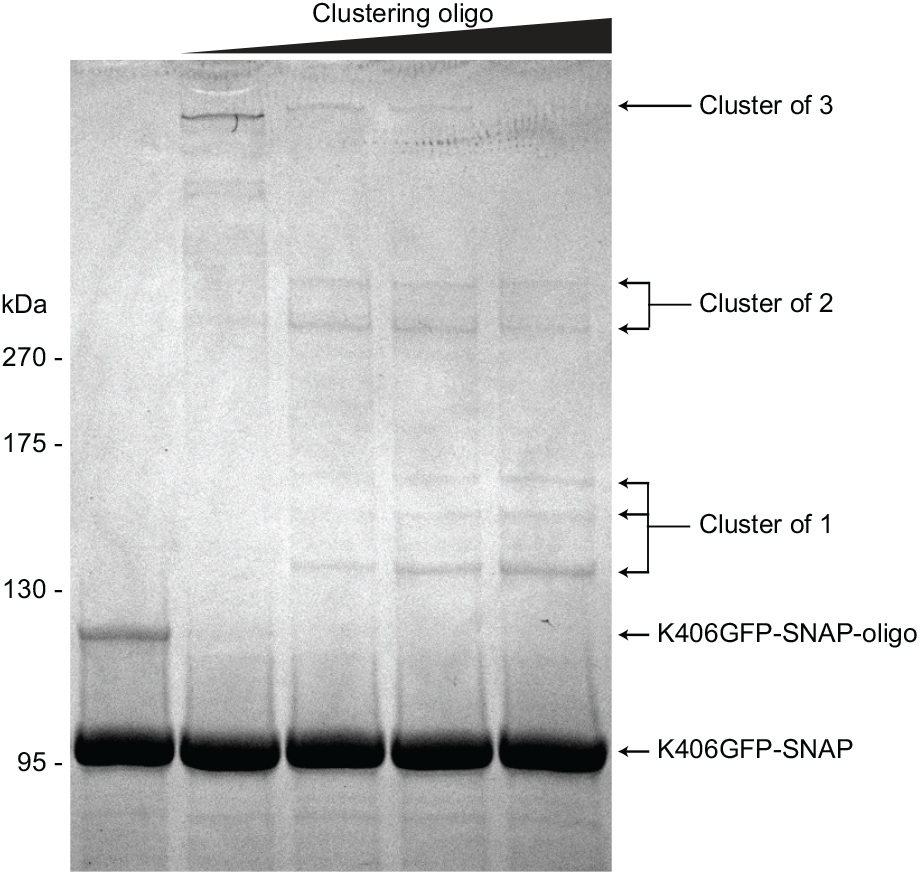
Electrophoretic mobility shift assay (EMSA) by SDS-PAGE confirms motor cluster formation. 15 fmol K406GFP-SNAP dimers mixed with increasing amount of the clustering oligo (left to right, 0, 2, 3.75, 5, 7.5 fmol) are loaded to each well. Proteins are stained by Coomassie blue.

**Supplementary Figure 5.**
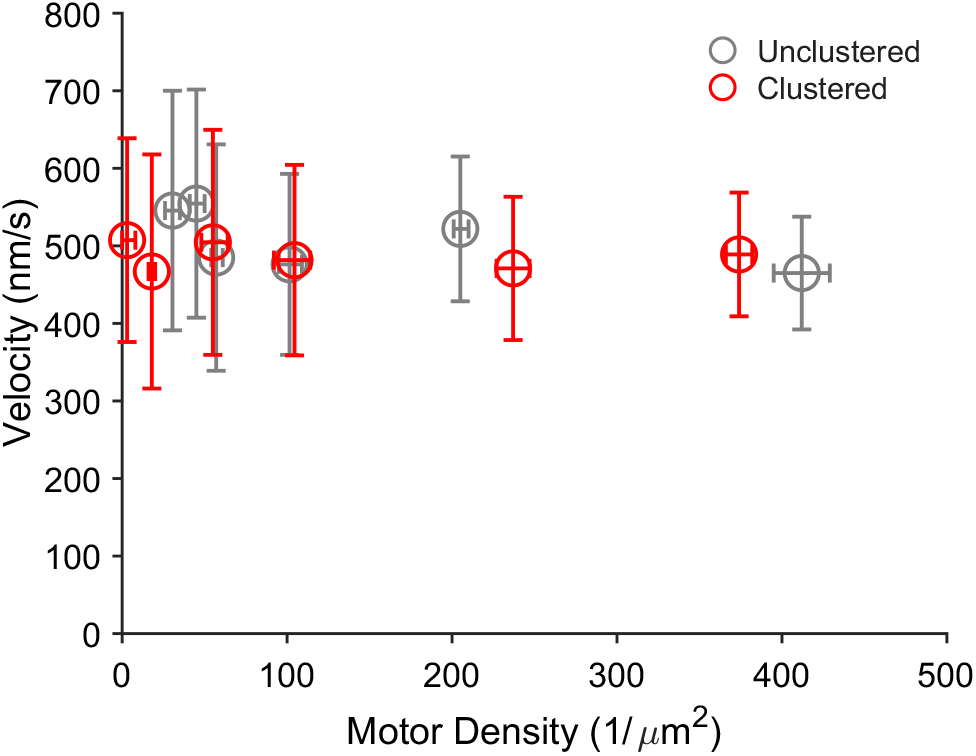
Liposome velocity at varying motor densities when motors are unclustered (gray) or clustered (red). Vertical error bars represent SD, horizontal error bars are calculated based on 10% increase in summed squared residuals.

## Notes

### Competing Interest Statement

The authors have declared no competing interest.

## Reference

1. A. G. Hendricks et al., Motor coordination via a tug-of-war mechanism drives bidirectional vesicle transport. Curr Biol 20, 697–702 (2010).

2. A. R. Chaudhary, F. Berger, C. L. Berger, A. G. Hendricks, Tau directs intracellular trafficking by regulating the forces exerted by kinesin and dynein teams. Traffic 19, 111–121 (2018).

3. E. E. Zahavi et al., Combined kinesin-1 and kinesin-3 activity drives axonal trafficking of TrkB receptors in Rab6 carriers. Dev Cell 56, 494–508 e497 (2021).

4. K. Furuta et al., Measuring collective transport by defined numbers of processive and nonprocessive kinesin motors. Proc Natl Acad Sci U S A 110, 501–506 (2013).

5. C. Herold, C. Leduc, R. Stock, S. Diez, P. Schwille, Long-range transport of giant vesicles along microtubule networks. Chemphyschem 13, 1001–1006 (2012).

6. M. Vershinin, B. C. Carter, D. S. Razafsky, S. J. King, S. P. Gross, Multiple-motor based transport and its regulation by Tau. Proc Natl Acad Sci U S A 104, 87–92 (2007).

7. R. P. Erickson, Z. Jia, S. P. Gross, C. C. Yu, How molecular motors are arranged on a cargo is important for vesicular transport. PLoS Comput Biol 7, e1002032 (2011).

8. A. Rai et al., Dynein Clusters into Lipid Microdomains on Phagosomes to Drive Rapid Transport toward Lysosomes. Cell 164, 722–734 (2016).

9. J. Beeg et al., Transport of beads by several kinesin motors. Biophys J 94, 532–541 (2008).

10. R. Jiang et al., Microtubule binding kinetics of membrane-bound kinesin-1 predicts high motor copy numbers on intracellular cargo. Proc Natl Acad Sci U S A 116, 26564–26570 (2019).

11. Q. Feng, K. J. Mickolajczyk, G. Y. Chen, W. O. Hancock, Motor Reattachment Kinetics Play a Dominant Role in Multimotor-Driven Cargo Transport. Biophys J 114, 400–409 (2018).

12. A. R. Chaudhary et al., MAP7 regulates organelle transport by recruiting kinesin-1 to microtubules. J Biol Chem 294, 10160–10171 (2019).

13. P. J. Hooikaas et al., MAP7 family proteins regulate kinesin-1 recruitment and activation. J Cell Biol 218, 1298–1318 (2019).

14. M. Metivier et al., Dual control of Kinesin-1 recruitment to microtubules by Ensconsin in Drosophila neuroblasts and oocytes. Development 146 (2019).

15. N. A. Reed et al., Microtubule acetylation promotes kinesin-1 binding and transport. Curr Biol 16, 2166–2172 (2006).

16. V. Soppina, A. K. Rai, A. J. Ramaiya, P. Barak, R. Mallik, Tug-of-war between dissimilar teams of microtubule motors regulates transport and fission of endosomes. Proc Natl Acad Sci U S A 106, 19381–19386 (2009).

17. I. A. Kesisova, B. P. Robinson, E. T. Spiliotis, A septin GTPase scaffold of dynein-dynactin motors triggers retrograde lysosome transport. J Cell Biol 220 (2021).

18. L. Urnavicius et al., Cryo-EM shows how dynactin recruits two dyneins for faster movement. Nature 554, 202–206 (2018).

19. A. A. Kendrick et al., Hook3 is a scaffold for the opposite-polarity microtubule-based motors cytoplasmic dynein-1 and KIF1C. J Cell Biol 218, 2982–3001 (2019).

20. A. R. Fenton, T. A. Jongens, E. L. F. Holzbaur, Mitochondrial adaptor TRAK2 activates and functionally links opposing kinesin and dynein motors. Nat Commun 12, 4578 (2021).

21. F. A. Ali et al., KIF1C activates and extends dynein movement through the FHF cargo adaptor. bioRxiv, 2023.2010. 2026.564242 (2023).

22. D. R. Klopfenstein, M. Tomishige, N. Stuurman, R. D. Vale, Role of phosphatidylinositol(4,5)bisphosphate organization in membrane transport by the Unc104 kinesin motor. Cell 109, 347–358 (2002).

23. W. G. Hirst, C. Kiefer, M. K. Abdosamadi, E. Schaffer, S. Reber, In Vitro Reconstitution and Imaging of Microtubule Dynamics by Fluorescence and Label-free Microscopy. STAR Protoc 1, 100177 (2020).

24. D. Nong et al., Integrated multi-wavelength microscope combining TIRFM and IRM modalities for imaging cellulases and other processive enzymes. Biomed. Opt. Express 12, 3253–3264 (2021).

25. Q. Li, K. F. Tseng, S. J. King, W. Qiu, J. Xu, A fluid membrane enhances the velocity of cargo transport by small teams of kinesin-1. J Chem Phys 148, 123318 (2018).

26. S. R. Nelson, K. M. Trybus, D. M. Warshaw, Motor coupling through lipid membranes enhances transport velocities for ensembles of myosin Va. Proc Natl Acad Sci U S A 111, E3986–3995 (2014).

27. J. D. Knight, M. G. Lerner, J. G. Marcano-Velazquez, R. W. Pastor, J. J. Falke, Single molecule diffusion of membrane-bound proteins: window into lipid contacts and bilayer dynamics. Biophys J 99, 2879–2887 (2010).

28. M. Uppalapati, Y.-M. Huang, S. Shastry, T. N. Jackson, W. O. Hancock, “Microtubule motors in microfluidics” in Methods in Bioengineering: Microfabrication and Microfluidics, J. D. Zahn, Ed. (Artech House Publishers, Boston, MA, 2009), chap. 13, pp. 311–336.

29. J. Schindelin et al., Fiji: an open-source platform for biological-image analysis. Nature methods 9, 676 (2012).

30. J. S. Verdaasdonk, J. Lawrimore, K. Bloom, Determining absolute protein numbers by quantitative fluorescence microscopy. Methods Cell Biol 123, 347–365 (2014).

